# Structural Basis for the Rational Design of UAMC-1110-Based FAP Radiopharmaceuticals

**DOI:** 10.64898/2026.10.11.758356

**Authors:** Chao Zhang, Jona Gerhards, Lara Scharbert, Jens Cardinale, Frederik L. Giesel, Birgit Strodel

## Abstract

Radioligands targeting fibroblast activation protein alpha (FAP) are promising tools for cancer diagnosis and therapy. However, the molecular binding mode of clinically established UAMC-1110-derived ligands such as FAPI-46 remains insufficiently characterized. To simplify synthesis while preserving key binding properties, we designed two novel derivatives—FAP-DD-101 (short aminoalkyl linker) and FAP-DD-102 (extended aminohexanoic acid spacer)—and evaluated them alongside FAPI-46 using molecular docking and all-atom molecular dynamics (MD) simulations. Across all derivatives, we identified a highly stable, energetically favorable charge-anchored binding mode, in which the 2-cyano-4,4-difluoropyrrolidine head anchors in a negatively charged subregion formed by Glu202, Glu203, and Glu204, mimicking the binding architecture observed in the FAP–linagliptin complex. MD trajectories confirmed that FAPI-46 and FAP-DD-101 maintain exceptional stability in this pose, with several non-anchored FAPI-46 trajectories spontaneously evolving toward engagement with this electrostatic subregion. For FAP-DD-102, linker elongation slightly reduced pose robustness, yet initial charge-anchored states still yielded superior average binding energies and vector stability compared to non-anchored poses. Overall, these findings elucidate the key structural determinants of FAP recognition, establishing a rational framework for designing synthetically accessible FAPI radiotracers with retained binding stability.

## 1 Introduction

Fibroblast activation protein alpha (FAP), also known as seprase, is a transmembrane protease that is overexpressed in the cancer-associated fibroblasts within the tumor microenvironment of a broad range of malignancies.^1–4^ FAP monomers are known to be inactive and require dimerization for activation.^5^ Structurally, each subunit of a FAP homodimer consists of 760 residues and can be divided into three parts: a 734 residue extracellular domain, a 20 residue transmembrane domain, and a 6 residue cytoplasmic domain. ^5,6^ FAP contains an enzymatic substrate domain with a catalytic triad comprising residues SER624, ASP702, and HIS734,^1^ which constitutes the binding sites for FAP inhibitors.

Due to the low expression of FAP in normal healthy adult tissues and its overexpression within the tumor microenvironment, FAP has emerged as an attractive molecular target for the development of diagnostic and therapeutic radiopharmaceuticals in nuclear medicine.^7–9^ The clinical potential of FAP-targeted radiopharmaceuticals has already been demonstrated in FAP-directed PET imaging for numerous malignancies, and is further reflected by the increasing number of ongoing FAPI clinical trials.^6,10^ A broad class of FAPI ligands was developed based on the UAMC-1110 scaffold^11^ (Figure 1) as the core binding motif, giving rise to several clinically relevant derivatives.^12,13^ Within the UAMC-1110 scaffold, the pyrrolidine-based ring incorporates difluoro substitution and a nitrile warhead, both of which contribute to enhanced binding affinity and/or selectivity toward FAP.^11^ Building on the UAMC-1110 scaffold, FAPI-01 and FAPI-02 were developed.^13^ FAPI-02 was designed to combine with the DOTA chelator, enabling radiolabeling with clinically relevant metallic radionuclides such as ^68^Ga, ^90^Y, ^177^Lu, and ^225^Ac,^14,15^ thereby providing diagnostic or therapeutic potential. However, FAPI-02 revealed a short tumor retention time in preclinical trials.^13^ Consequently, FAPI ligands were further optimized to improve tumor retention, resulting in the development of several new FAPI derivatives (FAPI-03 to FAPI-15).^12^ These derivatives were obtained by varying linker architectures at distinct positions of the quinoline ring and by tuning substituents on the pyrrolidine moiety. ^4^ Afterward, an additional FAPI series was developed, among which FAPI-46 emerged as the most promising candidate owing to its high tumor-to-background ratios and rapid tumor accumulation.^4,8^ The efficacy of ^68^Ga-labeled FAPI-46 has been evaluated in patients with various malignancies, including cholangiocellular, breast, gastric, head-and-neck, pancreatic, and oropharynx cancers. ^68^Ga-FAPI-46 PET imaging demonstrated favorable tumor visualization across a broad tumor spectrum with reduced radiation exposure.^16^

**Figure 1:**
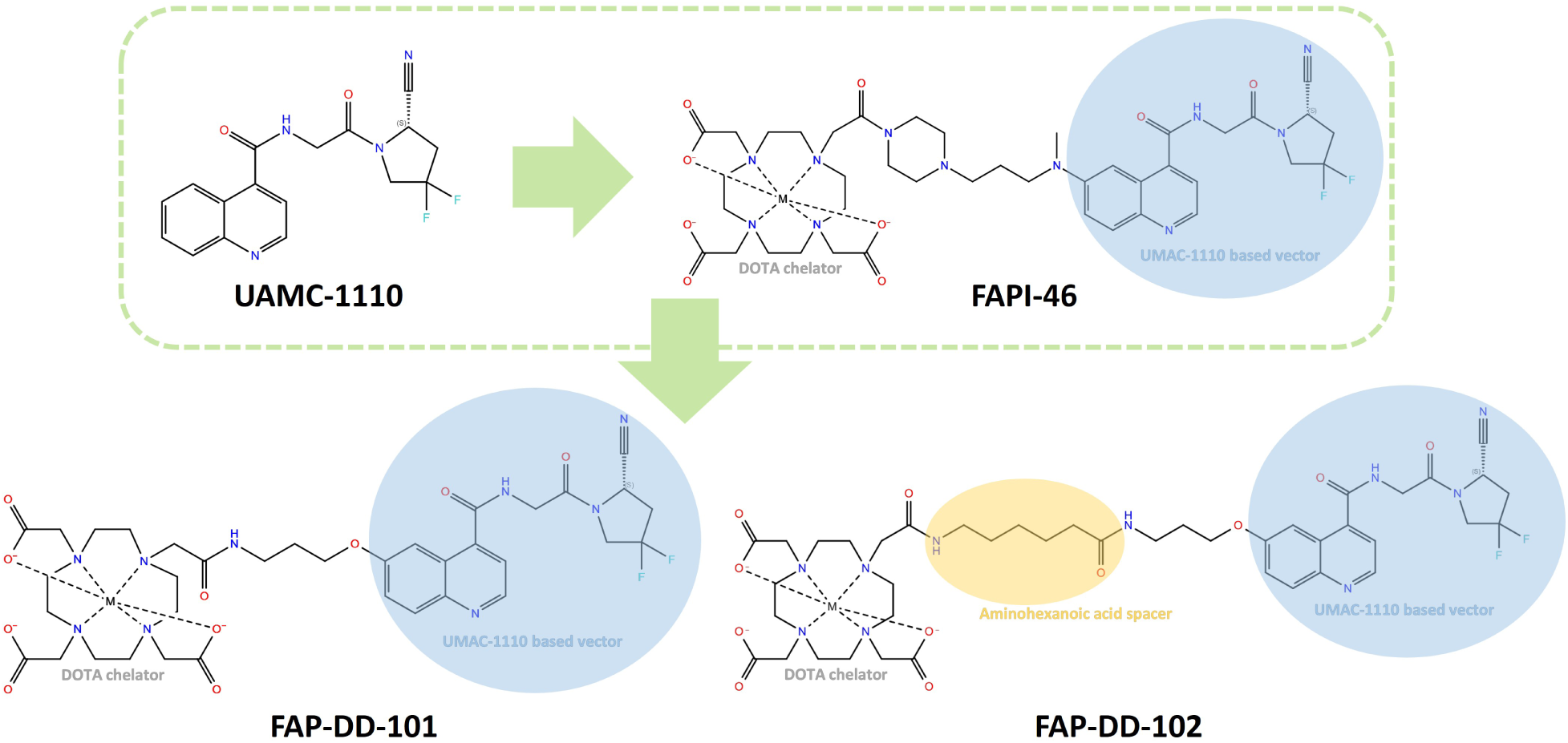
Structures of UAMC-1110, its derivative FAPI-46, and our newly designed ligands FAP-DD-101 and FAP-DD-102. The “M” within the chelator represents the radionuclide.

Although UAMC-1110–based FAPI radioligands have shown encouraging performance in tumor imaging and theranostic applications, the molecular binding mode underlying the interaction of these ligands with the FAP receptor remains insufficiently characterized. In particular, the preferred binding orientation of UAMC-1110–derived ligands within the FAP binding pocket and the relationship between binding orientation and conformational stability remain unclear.

To investigate these questions, FAPI-46 was selected as a representative UAMC-1110-derived ligand for detailed analysis. The piperazine-containing linker architecture of FAPI-46 remains synthetically demanding. To simplify ligand synthesis while maintaining favorable binding properties, we designed two novel UAMC-1110-based FAPI ligands: FAP-DD-101 and FAP-DD-102, using structurally simplified linker motifs (Figure 1). FAP-DD-101 was developed as an analog of FAPI-46 using a short aminoalkyl linker, whereas FAP-DD-102 incorporates an aminohexanoic acid spacer to extend the linker length. We sought to investigate whether the introduction of an aminoalkyl linker enhances ligand binding affinity, and whether the increased linker length influences the binding stability and binding mode of UAMC-1110–based ligands within the FAP binding pockets.

In this work, molecular docking and all-atom explicit-solvent molecular dynamics (MD) simulations were employed to investigate ligand-protein interactions and compare different binding orientations of UAMC-1110–derived FAPI ligands within the FAP binding pocket. Structural dynamics and binding energetics were further analyzed to evaluate the relationship between ligand orientation, binding stability, and relative binding favorability, thereby providing structural insight to support the rational design of future FAPI radioligands.

## 2 Methods

### 2.1 Receptor and ligand preparation

The three-dimensional structure of the FAP homodimer was obtained from the RCSB Protein Data Bank (PDB ID: 6Y0F, chains C and D). The FAP homodimer was prepared using the Protein Preparation Wizard^17^ in Maestro 13.5 (Schrödinger Release 2023-1)^18^ with the OPLS 2025 force field^19^ and a pH of 7.4 *±* 2.0.^20^ For the ligands FAPI-46, FAP-DD-101, and FAP-DD-102, the vector and linker moieties were constructed using the 2D Sketcher and converted to three-dimensional structures within Maestro. The DOTA chelator was subsequently incorporated based on its crystal structure retrieved from the Cambridge Crystallographic Data Centre (CCDC).^21^ Hydrogen atoms were then added to complete the ligand structures. Energy minimization of the vector and linker regions was performed using the OPLS 2005 force field to relieve unfavorable contacts. Protonation states of the ligands were predicted using the Epik Classic module^22^ at pH 7.0 *±* 2.0.^20^

### 2.2 Molecular docking

AutoDock FR 1.0 (ADFR 1.0)^23,24^ was employed for molecular docking. A technical challenge in docking the radioligands arises from the presence of the radionuclide ^68^Ga^3+^, which is not explicitly supported by most available docking programs, including ADFR 1.0 employed in this study. To enable docking calculations, Fe^3+^ was therefore adopted as a substitute metal ion. This substitution is justified by the comparable physicochemical properties of Fe^3+^ and Ga^3+^, including their identical formal charge, and similar ionic radius, ionization potentials, and electron affinity values.^25,26^ Moreover, classical molecular mechanics does not explicitly account for nuclear properties, rendering isotopic differences irrelevant within the present computational framework.

Two docking grids were defined for the two subunits of the prepared FAP homodimer and were centered on the coordinates of linagliptin in the FAP–linagliptin crystal complex (PDB ID: 6Y0F)^20^ with a padding of 10 Å. For each docking calculation, 120 independent runs were performed, with each run comprising 8 million evaluations. To validate the docking protocol, redocking using the crystallographic structure of linagliptin from the FAP–linagliptin complex (PDB ID: 6Y0F) was first conducted after completing missing hydrogen atoms using Maestro. FAPI-46, FAP-DD-101, and FAP-DD-102 were docked independently into both subunits of the FAP homodimer using the corresponding grids. Each docking generated multiple binding poses, which were ranked according to the ADFR scoring function.^23,27^ For post-docking analysis of ligand–protein interactions, electrostatic potential surfaces (EPS) were calculated using the Adaptive Poisson–Boltzmann Solver (APBS) electrostatics plugin^28,29^ implemented in PyMOL 2.5.^30^ After docking, selected top-ranked poses were subjected to MD simulations, after Fe^3+^ was replaced with Ga^3+^, and force field parameters were generated.

### 2.3 Ligand parameterization

The MCPB.py^31^ tool of AmberTools 23^32^ was employed to build force field parameters for the metal site in Ga-labeled ligands using the bonded model approach. Quantum mechanical (QM) geometry optimization, force constant calculation, and Merz-Kollman restrained electrostatic potential (RESP) charge calculation were performed at the B3LYP/6-31G* level of theory using Gaussian 09.^33^ The Seminario method was applied to derive metal ion–related bond and angle parameters from the QM force constants obtained in the Hessian calculation.^31,34^ After that, AMBER topology and coordinate files were generated with the LEaP module from AmberTools and converted into GROMACS-compatible input files with ACPYPE 2022.6^35^

### 2.4 MD simulations

All MD simulations were conducted in triplicate format according to the following protocol:

#### Setup

System setup was carried out using GROMACS 2018.6.^36^ Protein topologies were generated using the Amber14SB force field^37^ with Parmbsc1 force field supplement.^38^ Ligand parameters obtained from the ligand parameterization procedure were incorporated accordingly. Each protein–ligand complex was solvated in a periodic cubic box with a minimum distance of 1.0 nm between the solute and the box boundary, using the TIP3P water model. ^39^ Na^+^ and Cl^−^ ions were used to neutralize the system and to achieve a physiological ionic strength of 0.15 M. Subsequent energy minimization, equilibration, and production MD simulations were performed using GROMACS 2021.4.^40^

#### Energy minimization

Energy minimization was performed using the steepest descent algorithm^41^ until the maximum force on any atom was below 1000 kJ mol*^−^*^1^ nm*^−^*^1^.

#### Equilibration

To equilibrate the systems under physiological conditions (310 K and 1.0 bar), sequential simulations in the NVT (constant number of particles, volume, and temperature) and NPT ensemble (constant number of particles, pressure, and temperature) were carried out. NVT equilibration was performed for 100 ps using the leap-frog integrator^42^ and the V-rescale thermostat.^43^ This was followed by a 1 ns NPT equilibration, employing the V-rescale thermostat, the Berendsen barostat,^44^ and the leap-frog integrator. For both equilibration simulations, a time step of 2 fs was used.

#### MD production run

Production MD simulations were conducted for 100 ns under NPT conditions at 310 K and 1.0 bar using the Nose-Hoover thermostat^45^ and the Parrinello-Rahman barostat.^46^ A time step of 2 fs was used throughout. Electrostatic interactions were treated using the smooth Particle Mesh Ewald (SPME) method^47^ with a real-space cutoff of 1.2 nm. Long-range dispersion corrections for energy and pressure were applied with 1.2 nm as the distance for the Lennard-Jones (LJ) cut-off. Bonds with hydrogen atoms were constrained using the LINear Constraint Solver (LINCS) algorithm.^48^ Following the production runs, binding enthalpy calculations were performed using gmx MMPBSA version 1.6.2.^49,50^

## 3 Results and discussion

To investigate the binding behavior of UAMC-1110–derived FAPI ligands within the FAP binding pocket, molecular docking and subsequent EPS analysis were first performed to identify potential binding orientations and interaction patterns. Representative docking poses were subsequently subjected to MD simulations to predict binding dynamics. Binding stability and affinity were further analyzed to evaluate the relationship between ligand binding mode and binding behavior.

### 3.1 Binding pocket of the FAP receptor

The binding pocket of FAP is defined based on the co-crystallized structure of the FAP-linagliptin complex (PDB ID: 6Y0F),^20^ shown in Figure 2A. Linagliptin is known to bind within the catalytic region of FAP and inhibit its enzymatic activity, ^51,52^ thereby providing a structurally resolved reference for locating the binding site. This structure was selected to define the docking region, given the absence of available crystal structures of UAMC-1110–derived FAPI-ligands with FAP. We define the binding pocket in each FAP subunit as all residues within 6 Å of the crystallographic linagliptin, supplemented by ASP702 from the catalytic triad^1^ (Figure 2A). The 6 Å cutoff is selected to capture the major interaction residues while remaining consistent with commonly used distance-based criteria for protein–ligand interaction analysis.^53^ The inclusion of ASP702 ensures that the complete catalytic triad—SER624, ASP702 and HIS734—is represented in the pocket definition.

**Figure 2:**
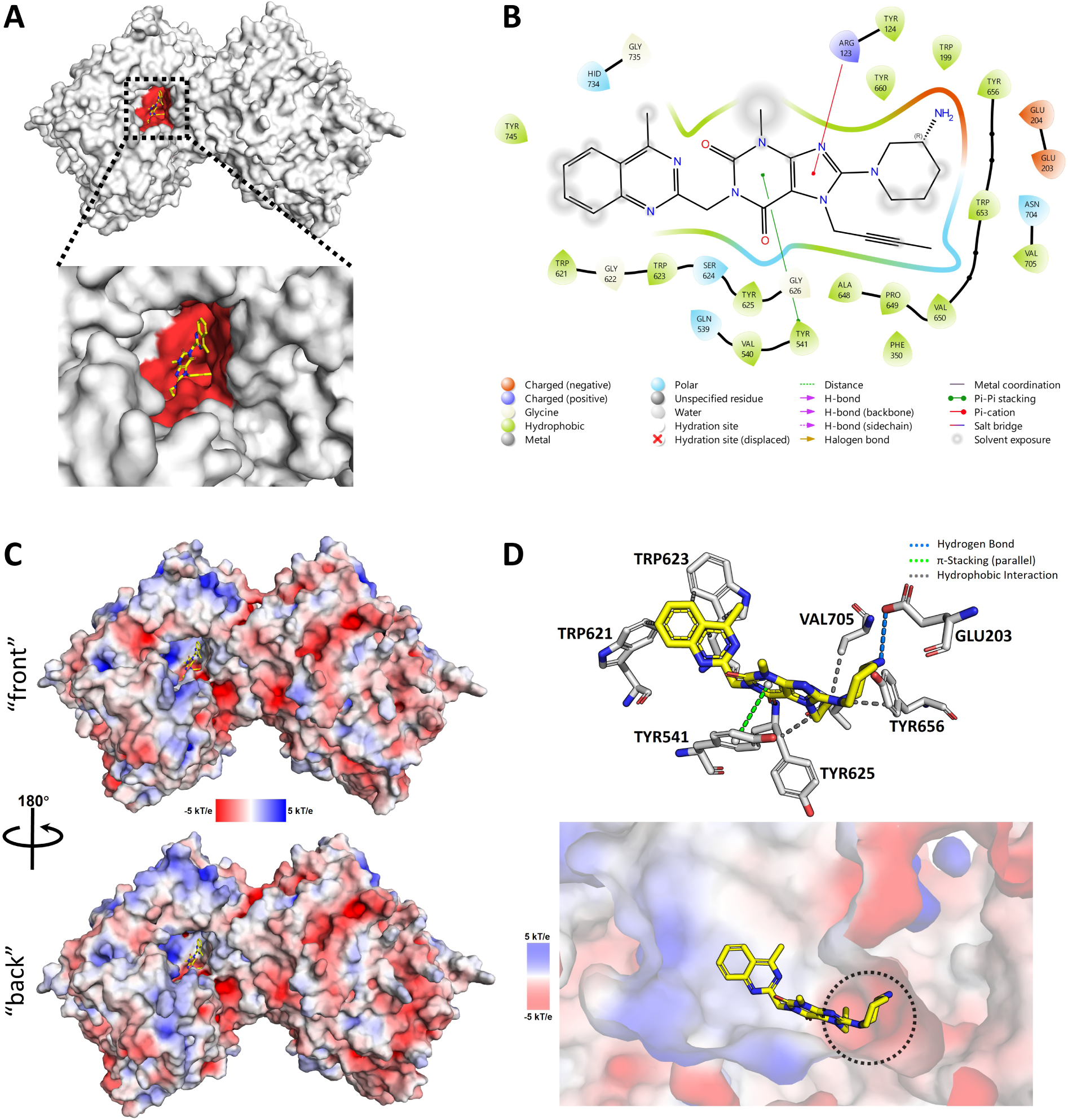
Structural and electrostatic characterization of the FAP binding pocket based on the FAP-linagliptin crystal structure (PDB ID: 6Y0F). (A) Binding pocket in one subunit of the FAP homodimer highlighted in red, defined as residues within 6 Å of the crystallographic linagliptin and supplemented by ASP702 from the catalytic triad. (B) Interaction diagram illustrating contacts between linagliptin and residues within the defined pocket. (C) Electrostatic potential surface of the FAP homodimer. The negatively charged regions are colored in red, and the positively charged regions are in blue. (D) Structural representation of protein-ligand interaction (top)^55^ and electrostatic representation of the binding pocket (bottom). The dotted circle emphasizes the distinct negatively charged binding subregion within the binding pocket.

The interaction map of the FAP-linagliptin complex (Figure 2B) indicates direct contacts between linagliptin and SER624 as well as HIS734, while residues adjacent to ASP702 (e.g., ASN704 and VAL705) are also positioned in the immediate vicinity of the ligand. Notably, GLU203 and GLU204 contribute to a negatively charged region within the binding pocket.

Together, these structural features indicate that the defined pocket (Figure 2A) corresponds to a ligand-accessible region, ^52,54^ and thus is subsequently used as the spatial reference for identifying and comparing binding modes of UAMC-1110–based FAPI ligands in docking and MD simulations.

To further elucidate the electrostatic environment governing ligand recognition, the EPS of the FAP homodimer was computed (Figure 2C). The two binding pockets of the homodimeric FAP display nearly identical electrostatic features. Therefore, only the binding pocket in subunit 1 is presented in detail. Despite the predominantly hydrophobic nature of the binding pocket, the EPS in Figure 2D reveals a heterogeneous charge distribution, characterized by distinct negatively and positively charged regions. Notably, a pronounced negatively charged subregion is observed near GLU203, as highlighted by the dotted circle in Figure 2D. This region primarily results from three glutamic acid residues, GLU202, GLU203, and GLU204, which collectively generate a localized negative electrostatic potential.

### 3.2 Docking pattern of FAPI-46, FAP-DD-101, and FAP-DD-102 with FAP receptor

After validating the reliability of the docking protocol and parameters via redocking, which reproduced the ligand crystal binding pose with a root-mean-square deviation below 1 Å (Figure S2), molecular docking was performed for FAPI-46, FAP-DD-101, and FAP-DD-102 against both binding pockets of the FAP homodimer, yielding multiple ranked binding conformations for each ligand. The binding poses were subsequently examined in the context of the EPS of the protein to evaluate their spatial and electrostatic complementarity. As each docking run generated multiple binding conformations with comparable docking scores, the top ten poses of each ligand were further analyzed. The top-ranked poses of the three ligands (FAPI-46, FAP-DD-101, and FAP-DD-102) exhibit comparable docking scores of around −10 kcal/mol (Figures 3, S3, and S4).

**Figure 3:**
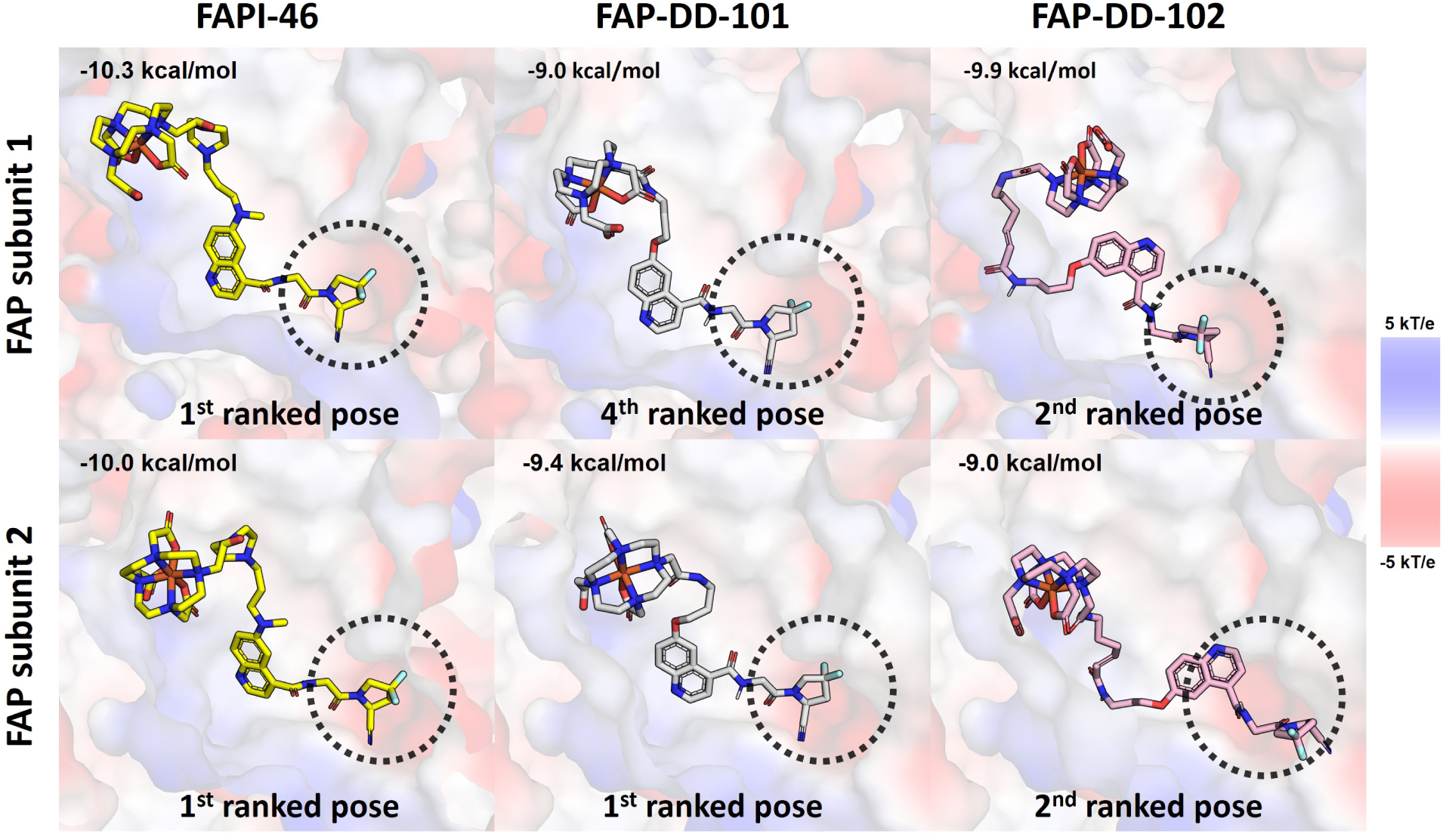
The highest-ranked ligand poses that follow the docking pattern in the two binding pockets of a FAP homodimer. The negatively charged regions are colored in red, and the positively charged regions are in blue. The negatively charged binding subregions are highlighted by dotted circles.

A consistent docking pattern is observed among the high-ranking poses of all three ligands (Figure 3). In this pattern, the 2-cyano-4,4-difluoropyrrolidine vector head is positioned within the negatively charged subregion highlighted by the dotted circle in Figure 3, while the chelator moiety extends toward slightly positively charged or neutral surface regions. Comparison between the docking pattern and the binding mode of linagliptin within the FAP pocket (Figure 2) reveals a common interaction feature, in which the pyrrolidine-based rings with polar groups are preferentially oriented toward the negatively charged subregions of the binding pockets. Previous studies have reported that the difluoro substitution and nitrile warhead of UAMC-1110 contribute to target binding affinity and/or selectivity.^11^ In the presented docking poses, the ring-containing head groups are consistently positioned near the same negatively charged subregions, suggesting that the local electrostatic environment may contribute to the observed binding orientation. In addition to the recurrent docking pattern, several high-ranking poses that do not follow this pattern were also identified (Figures S3 ans S4). These poses display diverse binding orientations while exhibiting docking scores comparable to those of the pattern poses. This illustrates a challenge associated with docking-based pose prediction, as scoring functions of docking may not always clearly distinguish between structurally distinct ligand orientations. Therefore, both pattern poses and representative non-pattern poses were selected for subsequent MD simulations to further evaluate their relative favorability.

### 3.3 MD simulations of FAP-ligand complexes

MD simulations offer a more physically rigorous description of ligand–protein interactions by explicitly accounting for atomic motions and solvent effects.^56,57^ Binding scores obtained from molecular docking rely on empirical scoring functions that enable rapid binding mode prediction, but employ simplified energetic representations of ligand–protein interactions.^23,58^ Moreover, docking inherently provides static snapshots of binding conformations and does not capture the dynamic behavior of complexes. To evaluate the dynamic stability of the proposed binding mode, all-atom explicit-solvent MD simulations were subsequently performed with ligand poses that follow the docking pattern (Figure 3), as well as for alternative poses that do not adopt this binding mode (Figure S4).

For each ligand—FAPI-46, FAP-DD-101, and FAP-DD-102—two different conformations were submitted to 100 ns MD simulations, each performed in triplicate. Both subunits of a FAP receptor contained a ligand that either initially followed the binding mode pattern (pattern pose) or adopted an alternative conformation (non-pattern pose). For each simulation, ligand dynamics and interaction behavior were characterized using structural descriptors, including conformational evolution, root-mean-square deviation (RMSD), root-mean-square fluctuation (RMSF), and binding energy profiles along the simulation trajectories.

#### 3.3.1 FAPI-46

FAPI-46, a clinically investigated UAMC-1110–derived FAPI ligand, was first selected as a representative ligand to examine whether the proposed docking pattern corresponds to a dynamically stable binding mode within the FAP binding pockets. By comparing pattern and non-pattern initial poses across independent MD simulations, we aim to assess the robustness of this binding orientation and its energetic favorability.

A comparative analysis of the pattern and non-pattern poses of FAPI-46 across the two binding pockets, derived from triplicate 100 ns MD simulations, is presented in Figure 4.

**Figure 4:**
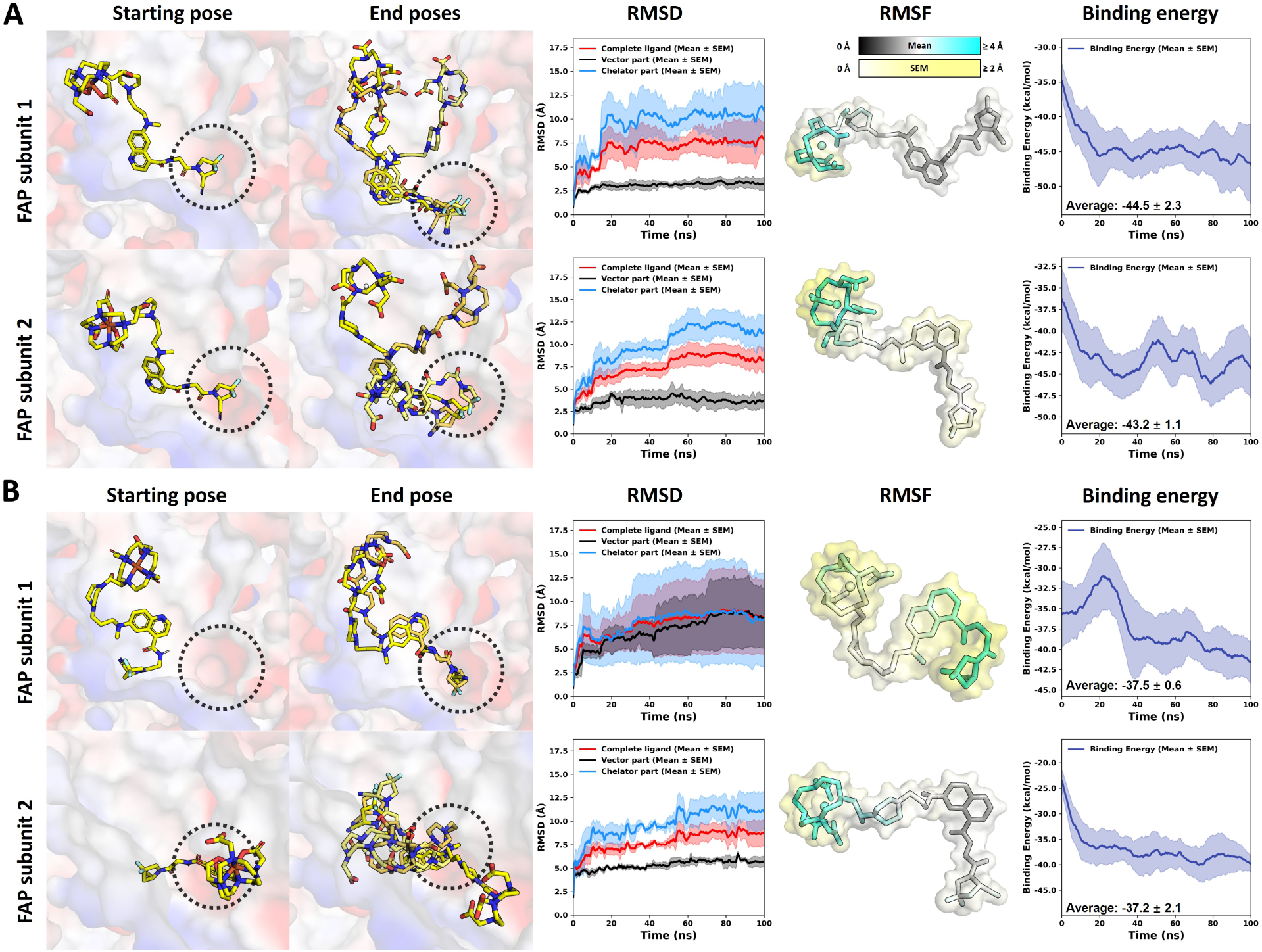
Structural analysis and binding energy evolution of FAPI-46 from triplicate MD simulations. (A) Pattern poses: initial ligand conformations that follow the proposed docking pattern in binding pockets 1 and 2 of the FAP homodimer. (B) Non-pattern poses: initial ligand conformations that do not follow the proposed docking pattern in binding pockets 1 and 2 of the FAP homodimer. For each system, three final ligand poses obtained from the three independent 100 ns MD simulations are shown. Only the binding regions of the protein are displayed as surfaces and colored according to their EPS, with negatively charged regions shown in red and positively charged regions shown in blue. The negatively charged binding subregions are highlighted by dotted circles. RMSD and binding energy profiles are reported as mean values with the standard error of the mean (SEM) from three independent simulations (solid lines with shaded areas). Mean ligand RMSF values from the triplicate simulations are mapped onto the ligand structures, while the corresponding SEM values are shown as surrounding color-coded surfaces; separate color bars above the structures indicate the scales for the mean RMSF and SEM values.

##### Pattern initial pose

Comparison of ligand conformations before and after the MD simulations (Figure 4A) shows that pattern poses preserve a consistent binding orientation of the vector head throughout each 100 ns trajectory. In all replicas, the pyrrolidine-based vector head remain anchored within the negatively charged binding subregion (dotted circles in Figure 4A), indicating a stable and persistent interaction. Consistent with this observation, RMSD profiles reveal limited structural deviation of the vector moiety relative to the chelator and the full ligand, with the vector regions exhibiting low RMSD values of around 2.5 Å in subunit 1 and around 3 Å in subunit 2. RMSF analysis further supports this stability, as the vectors display low flexibility (RMSF *≤* 2 Å). The chelator moieties show higher mobility, reflected by elevated RMSD *>* 8 Å and RMSF *≥* 4 Å. These results suggest that the negatively charged subregion contributes to the preferred orientation of the vector head, thereby promoting a stable binding mode, whereas the chelators do not provide interactions that enable the chelators to adopt a single conformation. Analysis of the binding energy evolution reveals progressive stabilization of the pattern poses over the course of the simulations, with binding energies decreasing as the ligands relax into more favorable conformations. The average binding energies of the pattern poses were approximately *−*44.5 and *−*43.2 kcal/mol in FAP subunits 1 and 2, respectively. Collectively, these findings demonstrate that ligand conformations in which the vector head occupies the negatively charged binding subregion correspond to stable binding modes.

##### Non-pattern initial poses

In the non-pattern initial pose replicas (Figure 4B), the vector heads migrate away from the starting configuration, suggesting that the non-pattern binding mode is not intrinsically stable. In particular, for the non-pattern initial poses in subunit 1, the ligand initially deviates from the proposed docking pattern by orienting the vector head toward a positively charged surface region rather than the negatively charged binding subregion. Remarkably, in two of the three MD replicas, the vector heads spontaneously migrate into the negatively charged binding subregion, resulting in a pronounced decrease in binding energy. Consequently, the binding energy decreases to approximately *−*42.3 kcal/mol toward the end of the simulations, approaching the values observed for the pattern poses. In contrast, the non-pattern pose in subunit 2 remained in non-pattern binding modes throughout the simulations, with binding energy remaining above *−*40.0 kcal/mol. Consistent with these observations, non-pattern initial poses exhibit larger average vector RMSD values, exceeding 5 Å in subunit 1 and approaching 5 Å in subunit 2. The vector heads in subunit 1 also exhibit RMSF values of *≥* 4 Å. These findings indicate that occupation of the negatively charged binding subregion is associated with a more energetically favorable and conformationally stable binding mode. Furthermore, the spontaneous transition from a non-pattern to a pattern binding mode highlights the ability of MD simulations to capture ligand relaxation and binding mode rearrangements beyond static docking poses.

#### 3.3.2 FAP-DD-101 and FAP-DD-102

To examine whether the stability of the proposed docking pattern is preserved across structurally related UAMC-1110–based FAPI ligands, we investigated our newly designed FAP-DD-101 and FAP-DD-102 using the same MD approach, which are summarized in Figures 5 and 6.

**Figure 5:**
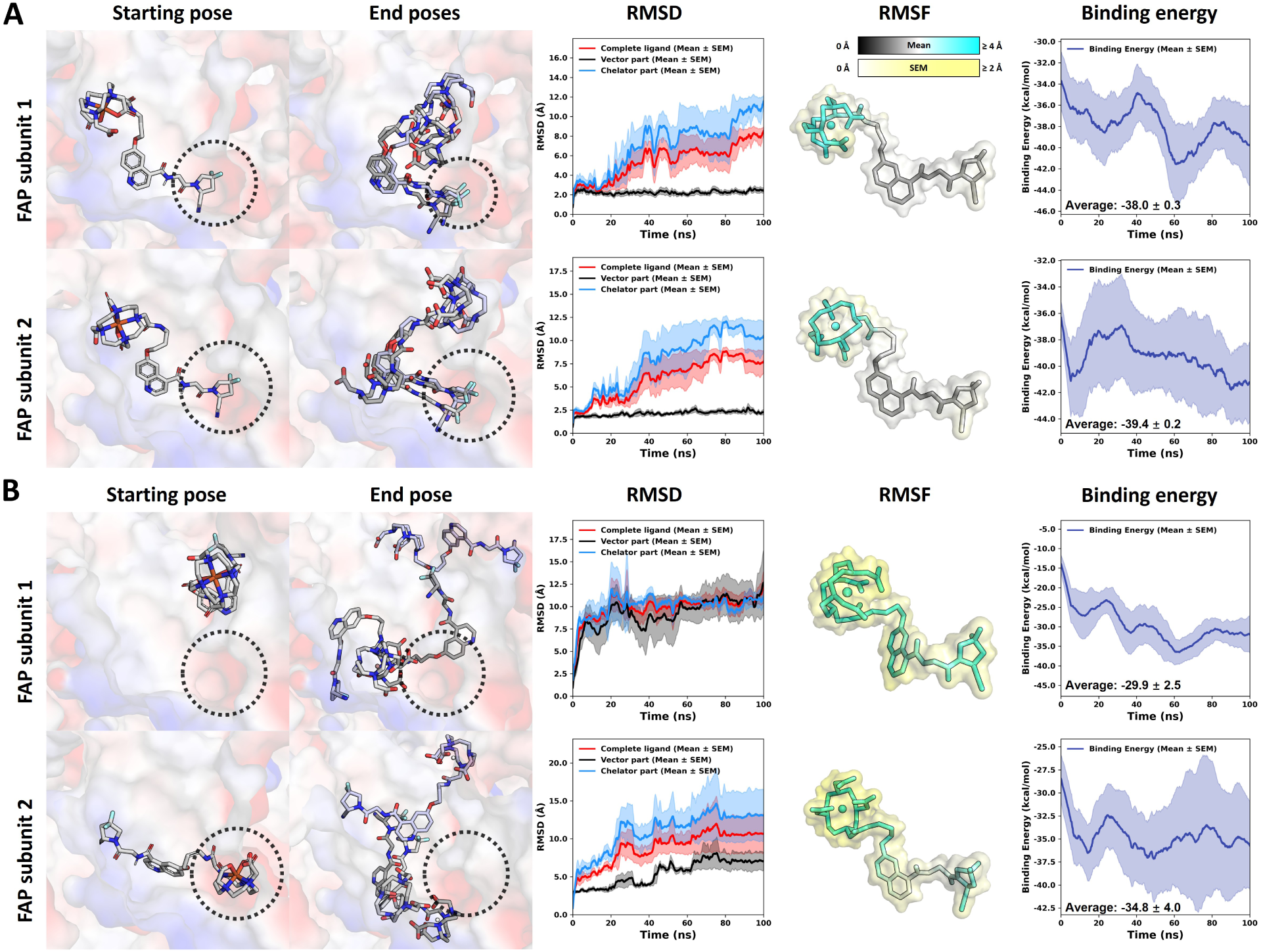
Structural analysis and binding energy evolution of FAP-DD-101 from triplicate MD simulations. (A) Pattern poses: initial ligand conformations that follow the proposed docking pattern in binding pockets 1 and 2 of the FAP homodimer. (B) Non-pattern poses: initial ligand conformations that do not follow the proposed docking pattern in binding pockets 1 and 2 of the FAP homodimer. For each system, three final ligand poses obtained from the three independent 100 ns MD simulations are shown. Only the binding regions of the protein are displayed as surfaces and colored according to their EPS, with negatively charged regions shown in red and positively charged regions shown in blue. The negatively charged binding subregions are highlighted by dotted circles. RMSD and binding energy profiles are reported as mean values with the standard error of the mean (SEM) from three independent simulations (solid lines with shaded areas). Mean ligand RMSF values from the triplicate simulations are mapped onto the ligand structures, while the corresponding SEM values are shown as surrounding color-coded surfaces; separate color bars above the structures indicate the scales for the mean RMSF and SEM values.

**Figure 6:**
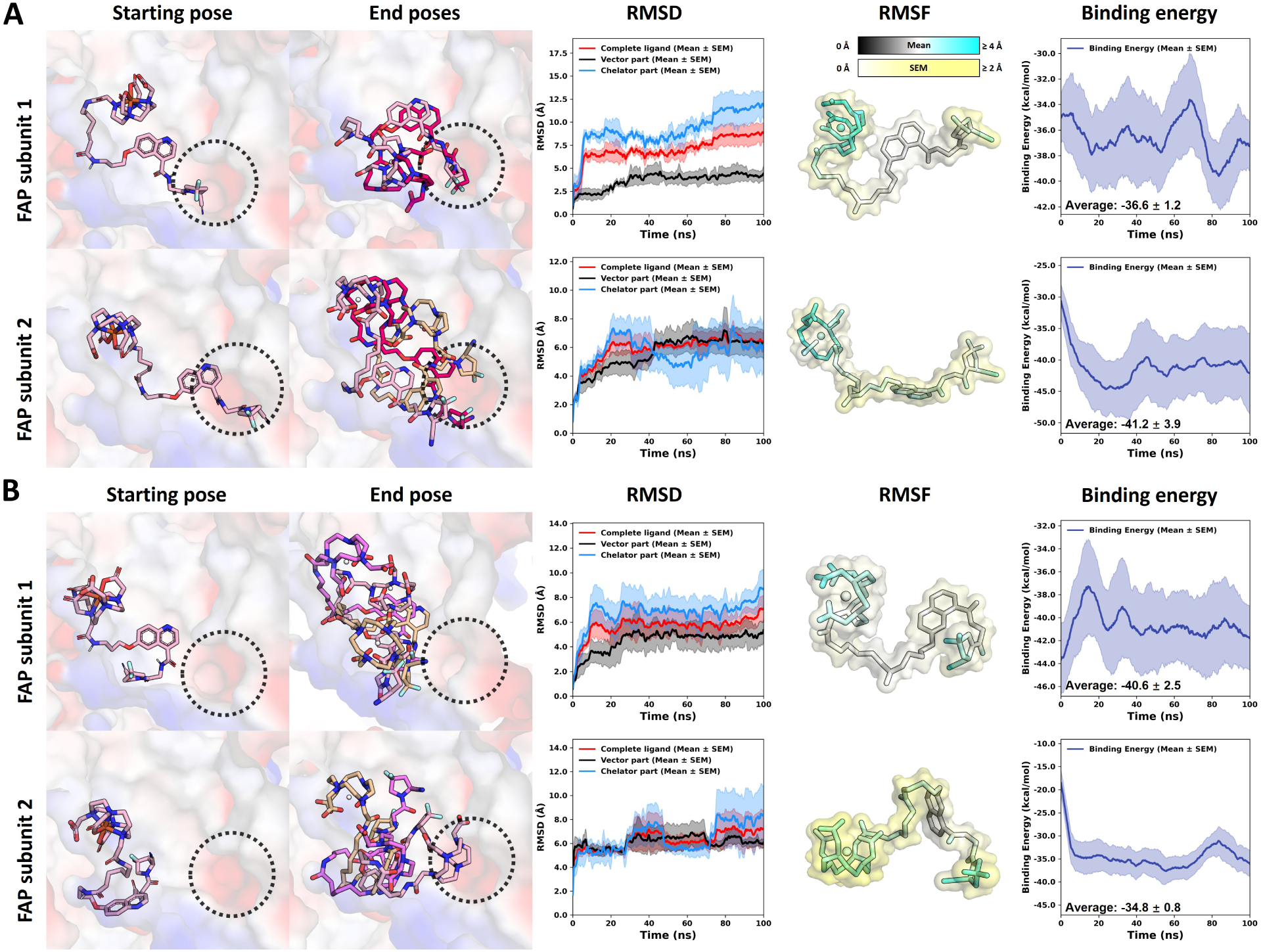
Structural analysis and binding energy evolution of FAP-DD-102 from triplicate MD simulations. (A) Pattern poses: initial ligand conformations that follow the proposed docking pattern in binding pockets 1 and 2 of the FAP homodimer. (B) Non-pattern poses: initial ligand conformations that do not follow the proposed docking pattern in binding pockets 1 and 2 of the FAP homodimer. For each system, three final ligand poses obtained from the three independent 100 ns MD simulations are shown. Only the binding regions of the protein are displayed as surfaces and colored according to their EPS, with negatively charged regions shown in red and positively charged regions shown in blue. The negatively charged binding subregions are highlighted by dotted circles. RMSD and binding energy profiles are reported as mean values with the standard error of the mean (SEM) from three independent simulations (solid lines with shaded areas). Mean ligand RMSF values from the triplicate simulations are mapped onto the ligand structures, while the corresponding SEM values are shown as surrounding color-coded surfaces; separate color bars above the structures indicate the scales for the mean RMSF and SEM values.

Similar to FAPI-46, FAP-DD-101 exhibits a stable association between the vector head and the negatively charged binding subregion. For ligands initiating from pattern poses, the vector head consistently remains positioned within this subregion throughout the simulations, exhibiting RMSD values below 2.5 Å and RMSF values below 2.0 Å. Furthermore, the linker and chelator regions display comparatively larger fluctuations (RMSF *≥* 2 Å), suggesting that these ligand segments are not constrained to a single conformation. Meanwhile, ligands initiating from non-pattern poses display heterogeneous vector conformations and less well-defined binding orientations at the end of the 100 ns simulations, indicating insufficient stabilization of the starting poses. The vector regions of these non-pattern ligands exhibit larger RMSD values of *>* 3 Å and elevated RMSF values of *≥* 2 Å, reflecting increased flexibility when not engaged within the proposed docking pattern. Consistent with the structural analyses, pattern initial poses of FAP-DD-101 yield lower binding energies throughout the simulations, with average values of approximately *−*38.0 and *−*39.4 kcal/mol in FAP subunits 1 and 2, respectively. These energies are consistently several kcal/mol lower than those of the non-pattern poses, further supporting that engagement of the vector head within the negatively charged binding subregion is associated with enhanced binding stability.

Unlike FAPI-46 and FAP-DD-101, FAP-DD-102 exhibits increased conformational variability during the MD simulations. For pattern initiating poses in FAP subunit 1, stable engagement of the vector head within the negatively charged binding subregion is maintained in only two of the three replicas, resulting in a worse average binding energy (−36.6 kcal/mol) compared with the pattern initiating poses in subunit 2 (−41.2 kcal/mol), where the vector heads remain in the negatively charged subregion in all three replicas. Nonetheless, the pattern initiating poses in FAP subunit 2 display reduced stability relative to FAPI-46 and FAP-DD-101, with a slight vector head deviation from the negatively charged binding subregion. Consistent with these trends, RMSD and RMSF analyses reveal increased flexibility of the vector head for FAP-DD-102 compared with FAP-DD-101. For pattern initial poses, the average vector RMSD values exceed 2.5 Å in subunit 1, and 4 Å in subunit 2, while the parts of the vectors exhibit RMSF values of *≥* 2 Å. This tendency become even more pronounced for non-pattern initial poses, with average vector RMSD values exceeding 4 Å in subunit 1 and approaching 6 Å in subunit 2, accompanied by RMSF values of *≥* 2 Å in portions of the vector head. These findings indicate that the vector head of FAP-DD-102 is less effectively stabilized within the negatively charged binding subregion than that of FAP-DD-101.

Given that FAP-DD-102 differs from FAP-DD-101 primarily by the incorporation of an aminohexanoic acid spacer, these observations indicate that the extended and more flexible linker introduces additional rotatable bonds, thereby enhancing conformational freedom and dynamic fluctuations within the binding pockets. Nevertheless, engagement of the vector head within the negatively charged binding subregion is still observed in multiple simulations, indicating that this region remains a favorable anchoring site despite the enhanced flexibility of FAP-DD-102. Overall, incorporation of the aminohexanoic acid spacer does not lead to a significant improvement in binding affinity, while increasing the propensity for deviation from the proposed binding mode.

### 3.4 Summary of docking and MD binding behavior across ligands

To provide an overview of the relationship between the docking results and the subsequent MD behavior, docking scores and average binding energies are summarized for FAPI-46, FAP-DD-101, and FAP-DD-102 in Table 1.

**Table 1:**
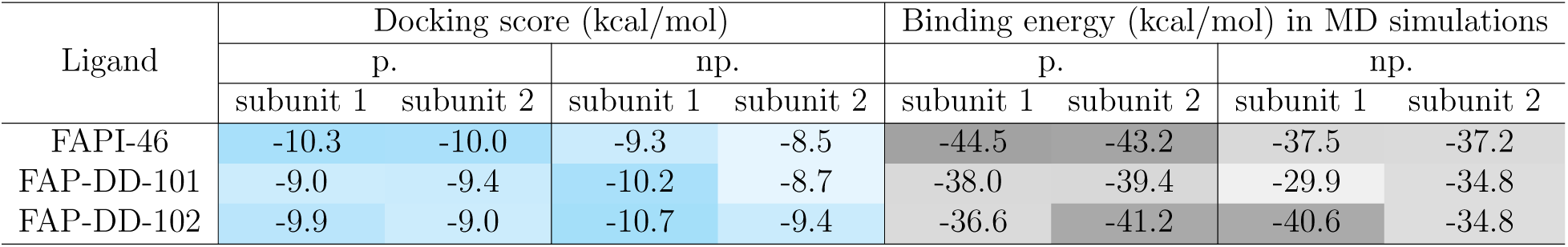
Binding affinities from docking and MD simulations. The p. stands for the highest-ranked poses following the pattern in docking and pattern initial poses in MD simulations. The np. stands for the highest-ranked poses that do not follow the pattern in docking and non-pattern initial poses in MD simulations.

In the docking results, FAP-DD-102 exhibits one non-pattern pose with a score of *−*10.7 kcal/mol, which is the most favorable docking score among all the poses of the three ligands. Additionally, FAPI-46 shows favorable scores for the two pattern poses, with docking scores of *−*10.3 and *−*10.0 kcal/mol.

Across the MD simulations, FAPI-46 displays the most favorable and stable binding behavior. For the initial pattern poses, FAPI-46 reaches average binding energies of *−*44.5 and *−*43.2 kcal/mol in the two FAP subunits, respectively, while maintaining the characteristic binding pattern. FAP-DD-102 also shows favorable binding in subunit 2, with an average binding energy of *−*41.2 kcal/mol. However, initial pattern poses in FAP subunit 1 show a weaker average binding energy of *−*36.6 kcal/mol, which is associated with the loss of the intended binding pattern in one replica, reflecting the higher flexibility and reduced stability of this ligand. Moreover, for all three ligands, the initial pattern poses always show more favorable average binding energies than the corresponding initial non-pattern poses.

Collectively, the MD simulations of FAPI-46, FAP-DD-101, and FAP-DD-102 demonstrate that engagement of the pyrrolidine-based vector head within the negatively charged binding subregion consistently corresponds to lower binding energies and enhanced structural stability. Ligands with shorter and less flexible linkers (FAPI-46 and FAP-DD-101) maintain this binding mode more robustly, whereas the introduction of the aminohexanoic acid spacer in FAP-DD-102 increases conformational freedom and dynamic fluctuations, partially weakening but not abolishing the preference for this binding subregion. These findings indicate that excessive linker flexibility introduces an entropic penalty that compromises binding stability without providing a significant compensatory gain in affinity, highlighting the importance of controlled linker length and rigidity in the rational design of UAMC-1110–based FAPI ligands.

## 4 Conclusion

In this study, molecular docking and MD simulations were used to investigate the binding behavior of UAMC-1110-based FAPI ligands in the FAP binding pocket. The combined simulations identified a potential stable binding mode for this ligand class, in which the 2-cyano-4,4-difluoropyrrolidine vector head preferentially anchors in a negatively charged subregion mainly formed by GLU202, GLU203, and GLU204. This binding mode is similar to that observed in the FAP-linagliptin complex crystal structure (PDB: 6Y0F),^20^ where a five-membered ring head bearing a polar amino substituent is positioned in the same negatively charged binding subregion, suggesting that this subregion may serve as a favorable recognition site for polar pyrrolidine-based vector heads, such as UAMC-1110.

For FAPI-46 and FAP-DD-101, this binding mode was dynamically stable and associated with favorable binding energies. Notably, several FAPI-46 trajectories initiated from non-pattern poses spontaneously evolved toward engagement of the negatively charged binding subregion, accompanied by improved binding energetics. For FAP-DD-102, although elongation of the linker by introducing an aminohexanoic acid spacer partially reduced the robustness of this binding mode, the pattern initial poses still showed more favorable average binding energies and greater vector stability than the non-pattern poses in MD simulations. Overall, identification and validation of this binding mode provide mechanistic insight into FAP recognition by UAMC-1110 derived radiopharmaceuticals and offer a rational structural framework for the future design of improved FAPI ligands.

## Acknowledgement

This research was funded by the Research Committee of the Medical Faculty (Düsseldorf, NRW, Germany), grant number FOKO 2023-39. The authors gratefully acknowledge computing time on the supercomputer JURECA at Forschungszentrum Jülich under grant “rdrp” as well as support by the Zentrum für Informations-und Medientechnologie (ZIM) at Hein-rich Heine University Düsseldorf for providing computational resources on the HPC cluster HILBERT. C. Zhang thanks his colleague Bastian F. Bundschuh for his helpful suggestions on the figures.

## Supporting Information Available

The following files are available free of charge.

- Structure of a human FAP homodimer in complex with linagliptin inhibitors.
- Redocking of linagliptin.
- The first-ranked ligand docking poses in the two subunits of a FAP homodimer.
- The highest-ranked ligand poses that do not follow the docking pattern in the two subunits of a FAP homodimer.
- Overview of end poses of ligands after triplicate 100 ns MD simulations.

## TOC Graphic

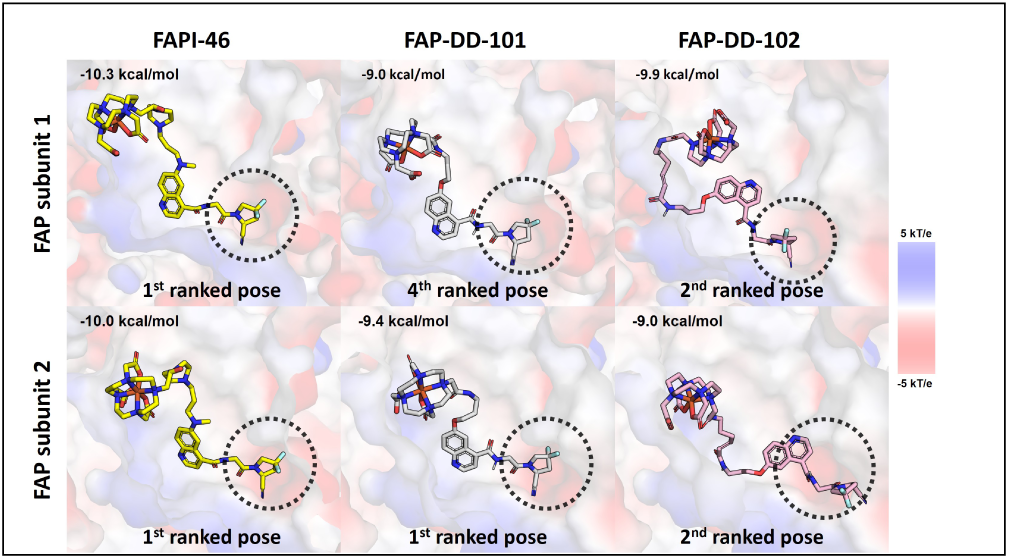

