## Supplemental Figures for "Structural Basis for the Rational Design of UAMC-1110-Based FAP Radiopharmaceuticals"

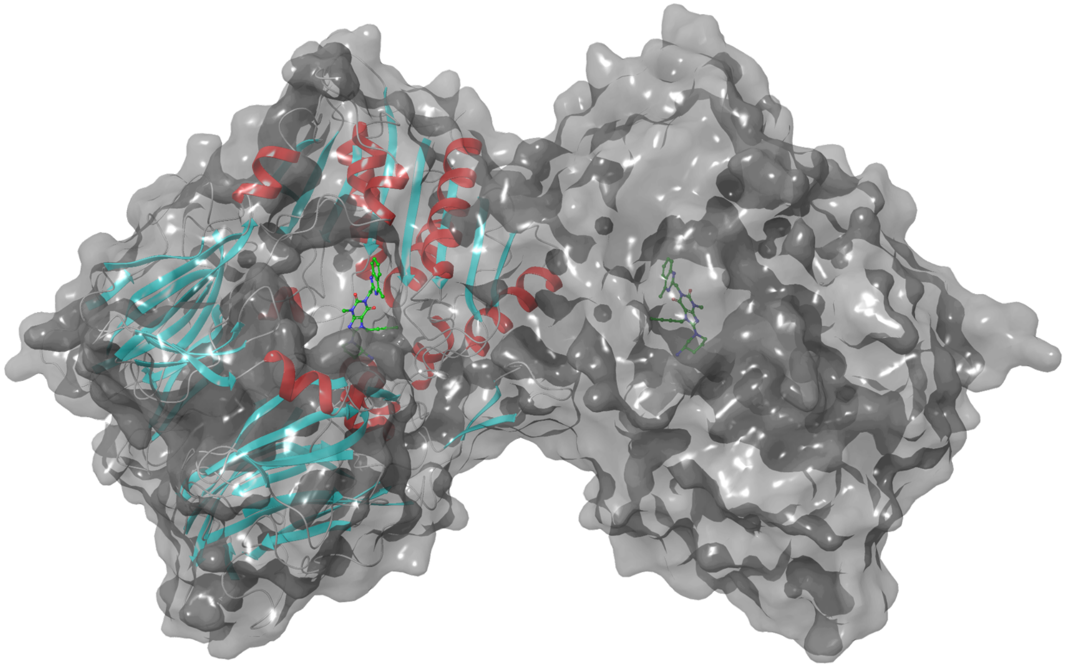

Figure S1: **Structure of human FAP homodimer in complex with linagliptin inhibitor.** The homodimer is represented by a molecular surface. One subunit of the dimer is also shown in ribbon cartoon style and colored based on the secondary structure. The inhibitor linagliptin is presented with green carbons. The figure was generated using Maestro software[1] and the protein structure is from the RCSB database (PDB ID: 6Y0F).[2]

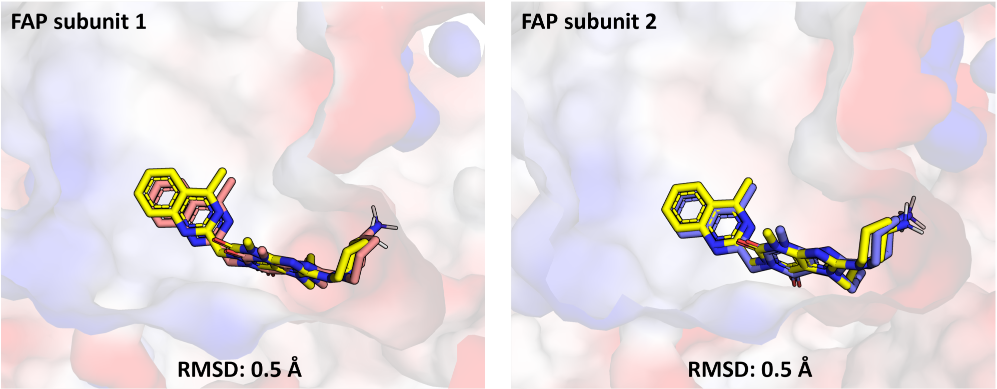


Figure S2: **Redocking of linagliptin.** The reliability of the docking protocol and parameter set was first assessed by redocking linagliptin into the FAP binding pocket prior to docking the ligands FAPI-46, FAP-DD-101, and FAP-DD-102. The docked poses of linagliptin closely reproduced the crystallographic conformation observed in the X-ray structure (PDB ID: 6Y0F), with root-mean-square deviation (RMSD) values below 1 Å. This high structural agreement confirms that the selected docking software and parameters are capable of generating experimentally relevant binding conformations. Here, the first-ranked redocking poses of linagliptin in both binding pockets of FAP dimer are shown, along with their RMSD values compared to the crystal structure.

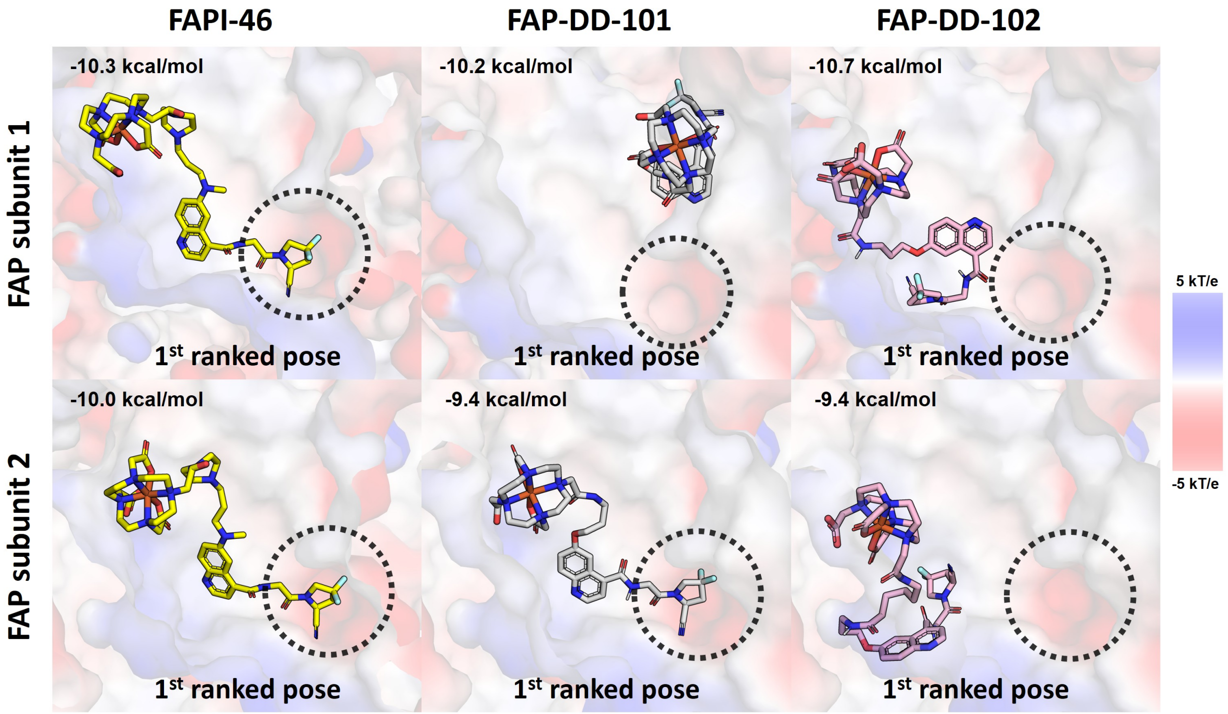


Figure S3: **The first-ranked ligand docking poses in the two subunits of a FAP homodimer.** The negatively charged regions are colored in red and the positively charged regions are in blue. The specific negatively charged binding subregions in the pockets are highlighted with dotted circles.

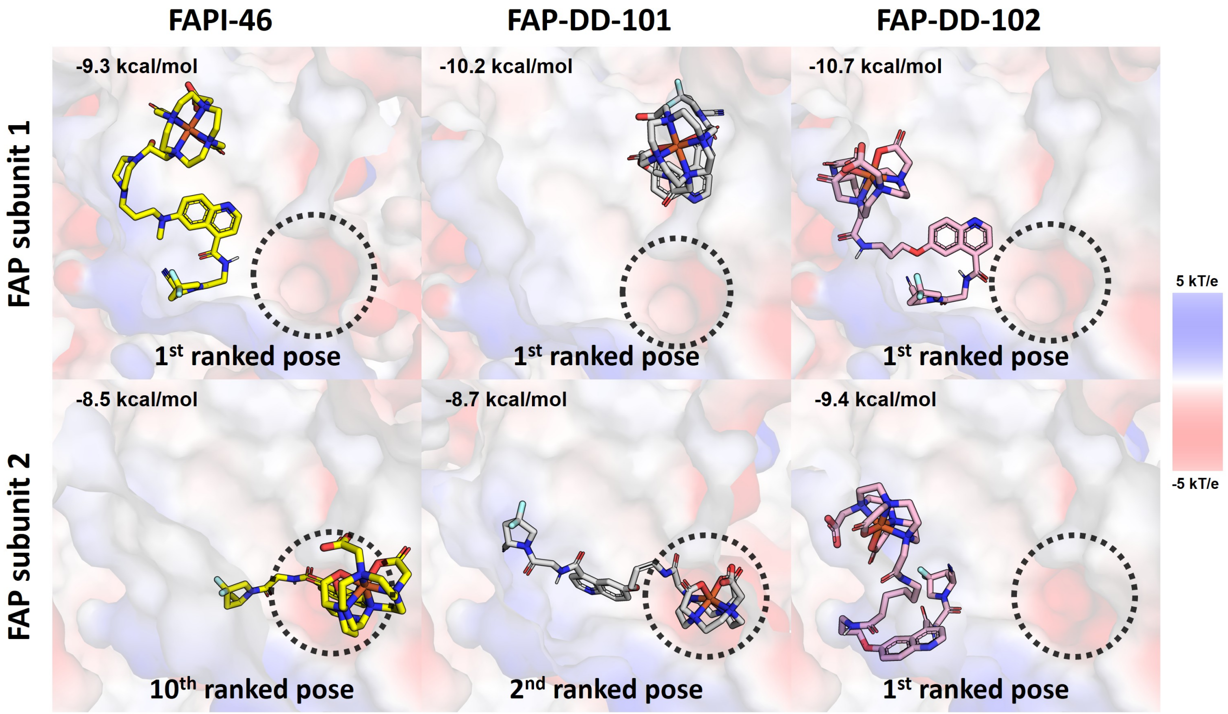


Figure S4: **The highest-ranked ligand poses that do not follow the docking pattern in the two subunits of a FAP homodimer.** The negatively charged regions are colored in red and the positively charged regions are in blue. The negatively charged binding subregions are highlighted by dotted circles.

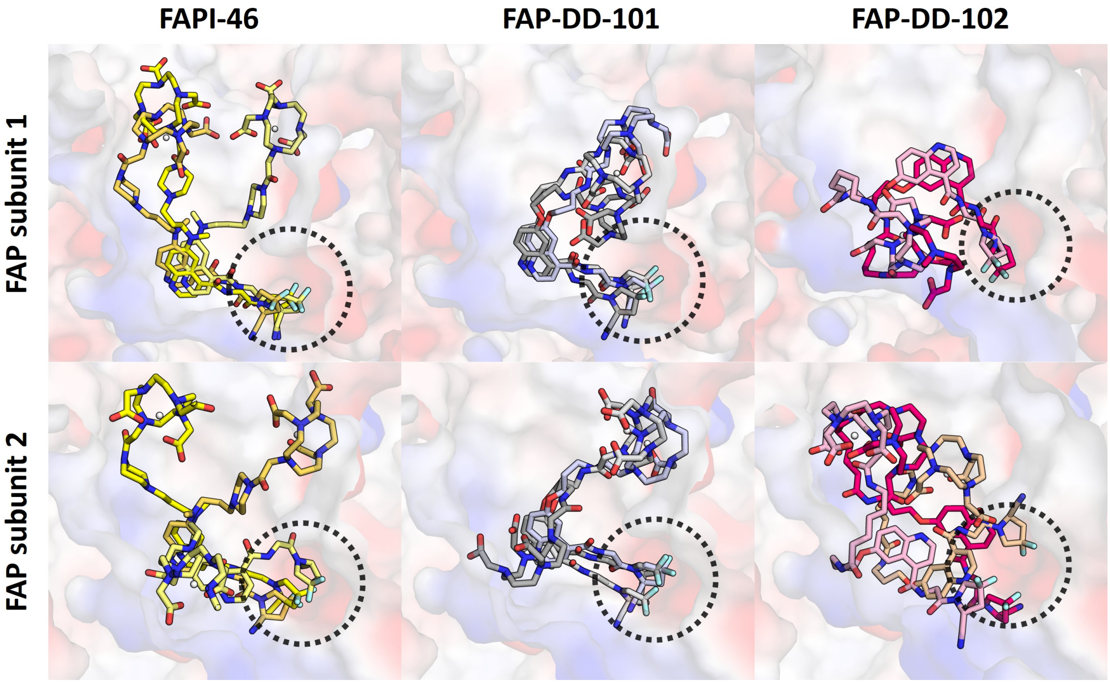


(**A**) End poses of ligands whose starting poses follow the pattern after triplicate 100 ns MD simulations.

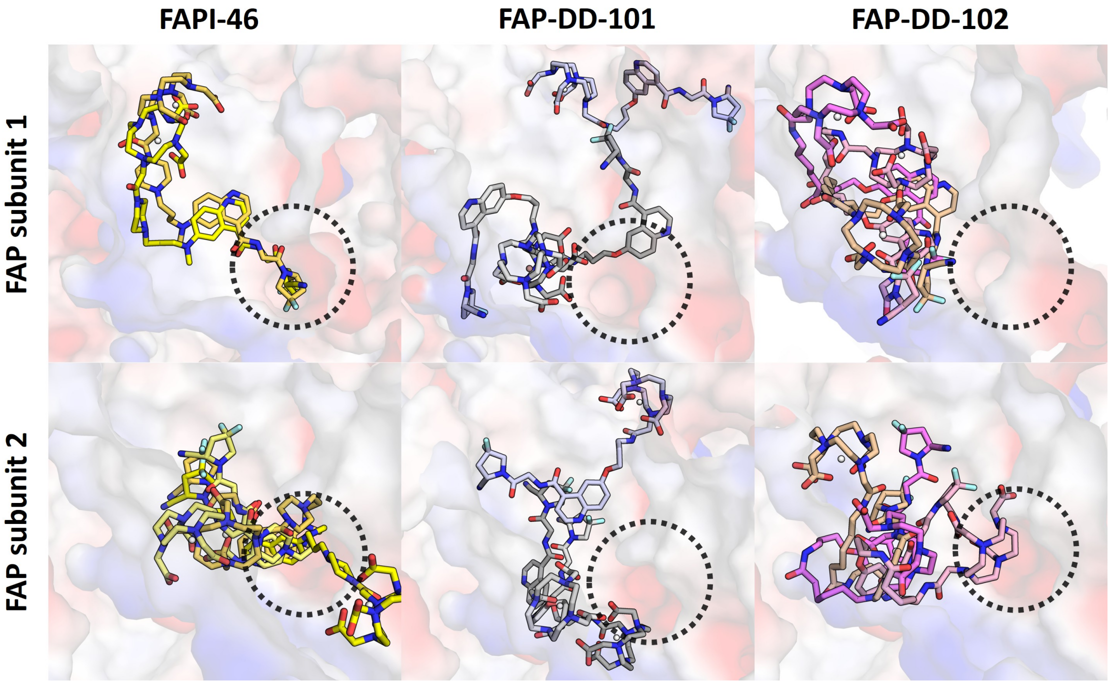


(**B**) End poses of ligands whose starting poses do not follow the pattern after triplicate 100 ns MD simulations.

Figure S5: **Overview of end poses of ligands after triplicate 100 ns MD simulations.** For each system, three final ligand poses obtained from the three independent 100 ns MD simulations are shown. The negatively charged regions are colored in red and the positively charged regions are in blue. The negatively charged binding subregions are highlighted by dotted circles.
